# Computational Analysis of Maize Enhancer Regulatory Elements Using ATAC-STARR-seq

**DOI:** 10.1101/2023.01.20.524917

**Authors:** Alexandre P. Marand

## Abstract

The blueprints to development, response to the environment, and cellular function are largely the manifestation of distinct gene expression programs controlled by the spatiotemporal activity of *cis*-regulatory elements. Although biochemical methods for identifying accessible chromatin – a hallmark of active *cis*-regulatory elements – have been developed, approaches capable of measuring and quantifying *cis*-regulatory activity are only beginning to be realized. Massively Parallel Reporter Assays coupled to chromatin accessibility profiling present a high-throughput solution for testing the transcription-activating capacity of millions of putatively regulatory DNA sequences in parallel.

However, clear computational pipelines for analyzing these high-throughput sequencing-based reporter assays are lacking. In this protocol, I layout and rationalize a computational framework for the processing and analysis of Assay for Transposase Accessible Chromatin profiling followed by Self-Transcribed Active Regulatory Region sequencing (ATAC-STARR-seq) data from a recent study in *Zea mays*. The approach described herein can be adapted to other sequencing-based reporter assays and is largely agnostic to the model organism with the appropriate input substitutions.

## [Background]

Eukaryotic cells exhibit remarkable functional and morphological diversity despite containing a generally invariant copy of the same genomic sequence. Cellular heterogeneity arises in part due to the activities of *cis*-regulatory elements (CREs), short DNA binding motifs recognized by sequence specific transcription factors (TFs). CREs are often found in clusters termed *cis*-regulatory modules (CRMs) that dictate highly dynamic spatiotemporal patterns of gene expression via the cooperative activities of DNA-bound TFs (Schmitz *et al*., 2022). For proper activation of transcription, the cell strictly regulates CRM activity by controlling TF access of CRM sequences through nucleosome dynamics. Genome-wide approaches, such as Assay for Transposase Accessible Chromatin sequencing (ATAC-seq), have been developed to profile accessible chromatin regions (ACRs) (Buenrostro *et al*., 2013; Minnoye *et al*., 2021). In general, CRMs that localize to accessible chromatin reflect active regulatory elements (Marand *et al*., 2017; Schmitz *et al*., 2022). Thus, activation and silencing of gene expression is effectively controlled by the relative chromatin accessibility of cognate CRMs.

CREs can be classified into distinct functional groups based on their regulatory effect on transcription, including enhancers, silencers, promoters, and insulators (Schmitz *et al*., 2022). Of these, enhancers are of particular interest due to their transcription activating properties that function independent of location and orientation of their target genes, in contrast to the stereotypical locations of promoters surrounding gene transcription start sites (TSSs) (Marand *et al*., 2017; Schmitz *et al*., 2022). While analysis of chromatin accessibility in distinct tissues and cell types has been central to identification of CRMs (Marand *et al*., 2021), chromatin profiling techniques are largely qualitative and lack the ability to quantitatively estimate regulatory activity. To overcome these challenges, Massively Parallel Reporter Assays (MPRA) have been developed to quantify the transcription activating properties of diverse sequences (Melnikov *et al*., 2012; Arnold *et al*., 2013;). In particular, Self-Transcribing Active Regulatory Region with sequencing (STARR-seq) demonstrates the greatest potential for broad application by eliminating the need for homogenous cell lines available only in mammalian models typical of other MPRA methods (Arnold *et al*., 2013; Ricci *et al*., 2019; Sun *et al*., 2019; Jores *et al*., 2020). Although STARR-seq was originally designed to profile the entire genome for regulatory activity, recent implementations have successfully utilized ATAC-seq libraries as input (ATAC-STARR-seq), reducing the search space to potential regulatory regions and offsetting sequencing costs and library complexity requirements (**Figure 1**). Despite its promise as a powerful approach towards understanding *cis*-regulatory activity, computational analysis of ATAC-STARR-seq data remains challenging, particularly due to a lack of dedicated software and computational pipelines.

**Figure 1.**
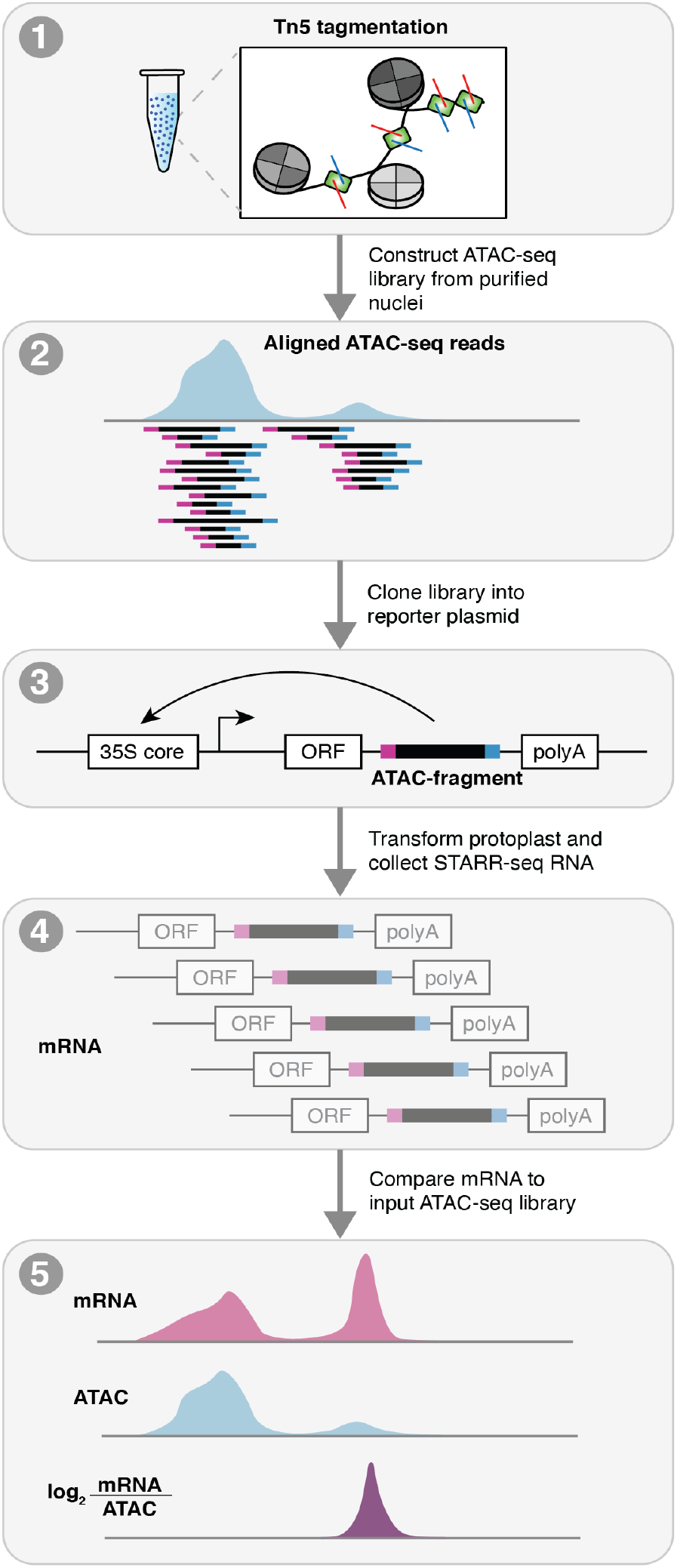
Schematic of ATAC-STARR-seq. ATAC-STARR-seq begins by first generating an ATAC-seq library. The ATAC fragments are then cloned into a reporter assay and transformed into maize protoplasts. After X hours, transformed protoplasts are then split into two pools, the first for sequencing the input fragments (ATAC-seq DNA), and the second for purifying transcribed (mRNA) ATAC-seq fragments that facilitate their own transcription from the reporter construct. Raw sequenced reads for the ATAC-seq input and mRNA output are processed and aligned to the maize reference genome and compared to provide estimates of cis-regulatory activity.

Here, I present a computational pipeline for analysis of ATAC-STARR-seq data generated in *Zea mays* L., cultivar B73 (Ricci *et al*., 2019). After processing and evaluation of data quality, I demonstrate how ATAC-STARR-seq data analysis allows for the interrogation of new biological questions. The pipeline can be run entirely from the code below or through freely available bash, perl, and R scripts hosted at https://github.com/Bio-protocol/Maize_ATAC_STARR_seq.

### Equipment

This pipeline assumes that a user has knowledge of shell commands and is comfortable working on a Linux-based operating system.

1. Computational Requirements

The following procedure can be run on any Linux-like system. However, this protocol and publicly available code is written for executing commands via a high-performance computing (HPC) cluster managed by a SLURM scheduler. However, the code presented here can be readily converted to TORQUE or other HPC systems. The pipeline assumes a working Perl interpreter version 5.30.0 or greater, and R version 3.6.2 or greater.

### Software

The following analytical procedure makes use of several standard computational tools that are assumed to be available in the user’s shell environment.

#### Software

1. *BWA MEM* (Li and Durbin, 2009); v0.7.17; http://bio-bwa.sourceforge.net/bwa.shtml
2. *SAMtools* (Li *et al*., 2009); v1.14; http://www.htslib.org
3. *BEDtools* (Quinlan and Hall, 2010); v2.27.1; https://bedtools.readthedocs.io/en/latest/
4. *SRA-toolkit* (Leinonen *et al*., 2011); v2.11.1; https://github.com/ncbi/sra-tools
5. *fastp* (Chen *et al*., 2018); v0.20.0; https://github.com/OpenGene/fastp
6. *pigz* v2.4; https://zlib.net/pigz/
7. *MACS2* (Liu, 2014); v2.2.7.1; https://pypi.org/project/MACS2/
8. *UCSC binaries* (Kent *et al*., 2010); v1.04.0; http://hgdownload.soe.ucsc.edu/admin/exe/linux.x86_64/
9. *tabix* (Li, 2011); v0.2.6; http://www.htslib.org/doc/tabix.html
10. *IGV* (Thorvaldsdottir *et al*., 2013); v2.11.1; https://software.broadinstitute.org/software/igv
11. *MEME* (Grant *et al*., 2011); v5.4.1; https://meme-suite.org/meme/index.html
12. *CrossMap* (Zhao *et al*., 2014); v0.5.1; http://crossmap.sourceforge.net
13. *DeepTools* (Ramirez *et al*., 2014); v3.5.1; https://deeptools.readthedocs.io/en/develop/index.html

#### Input data

The starting input for this computational pipeline uses paired-end sequencing data from an ATAC-STARR-seq experiment performed on maize protoplasts (Ricci *et al*., 2019). The ATAC-STARR-seq experiment consisted of a DNA input (ATAC-seq library) and a mRNA readout (self-transcribed regulatory regions) to identify genomic regions exhibiting transcription-activating regulatory activity.

1. Transfected ATAC-seq DNA-input FASTQ
2. Transcribed ATAC-seq mRNA FASTQ

### Procedure

An overview of the ATAC-STARR-seq pipeline is presented in **Figure 2**.

**Figure 2.**
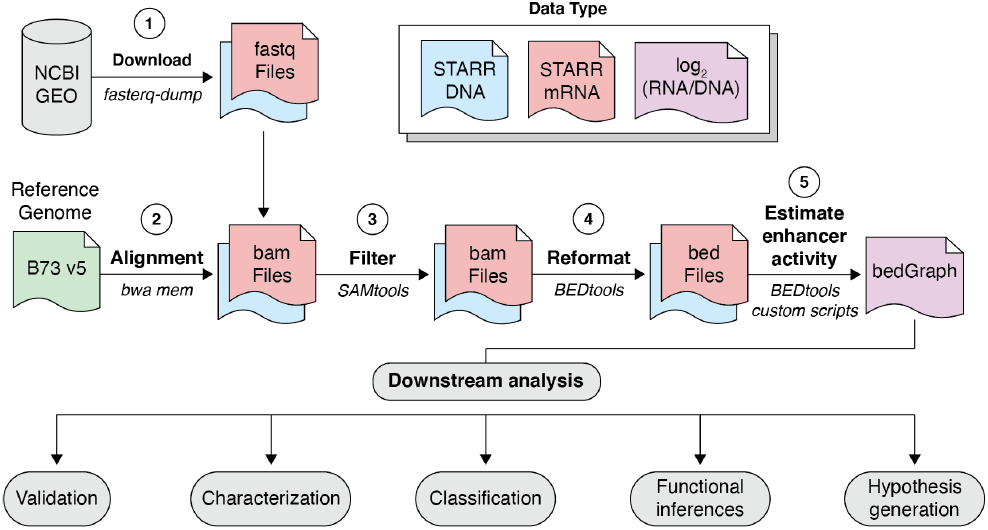
Computational workflow for ATAC-STARR-seq data. Raw sequence ATAC-STARR-seq data is first acquired from NCBI GEO, processed, and aligned to the reference genome with *bwa mem*, filtered via *samtools*, reformatted as fragments with *bedtools*, and compared via *bedtools* and custom *R* scripts to provide estimates of enhancer activity for downstream analysis.

#### A. Download and prepare the requisite data and reference genome sequence

1. Raw mRNA ATAC-STARR-seq data generated from transfection of *Zea mays* leaf ATAC-seq fragments in *Zea mays* protoplasts, and the accompanying ATAC-seq input fragments, are publicly available on NCBI GEO. ATAC-STARR-seq mRNA and DNA input can be downloaded with *fasterq-dump* available from the *SRA-Toolkit* package:

~~~
# set variables and download FASTQ files
mkdir FASTQ_files
cd FASTQ_files
fasterq-dump -o B73_maize_DNA_input.fastq SRR10964904
fasterq-dump -o B73_maize_mRNA_output.fastq SRR10964905
~~~
2. To save disk space, we will compress the FASTQ files with *pigz*. By default, *pigz* uses all available processors or eight if the number of available processors is unknown. Alternatively, users can use the unix tool, *gzip*, to compress the STARR mRNA and ATAC input DNA FASTQ files.

~~~
# compress fastq files
pigz *.fastq
# NOT RUN
# Tip: gzip can be used as an alternative to pigz (parallel gzip)
# gzip *.fastq
~~~
3. Download the B73 reference genome sequence and gene annotation. The original article mapped raw reads to version 4 of the B73 maize reference genome. In this case study, I map and analyze maize ATAC-STARR-seq data to version 5 of the B73 maize reference genome to showcase how updated reference genomes and read mapping strategies enable informative reanalysis of publicly available data sets (Hufford *et al*., 2021).

~~~
# download reference data
cd ../
mkdir Genome_Reference
cd Genome_Reference
wget https://download.maizegdb.org/Zm-B73-REFERENCE-NAM-5.0/Zm-B73-REFERENCE-NAM-5.0.fa.gz
wget https://download.maizegdb.org/Zm-B73-REFERENCE-NAM-5.0/Zm-B73-REFERENCE-NAM-5.0_Zm00001eb.1.gff3.gz
~~~

To map data to the genome reference, we first need to decompress the FASTA file. Constructing reference genome indices is a prerequisite for *BWA* alignment and allows for faster post-alignment processing of BAM/SAM/BED formatted files with command line tools such as *SAMtools* and *BEDtools*.

~~~
# create indices for reference genome FASTA
gunzip Zm-B73-REFERENCE-NAM-5.0.fa.gz
samtools faidx Zm-B73-REFERENCE-NAM-5.0.fa
bwa index Zm-B73-REFERENCE-NAM-5.0.fa
~~~

#### B. Trim adapters and remove low quality reads

1. Illumina platforms may produce reads with adapter sequences on the 3’ ends if the DNA insert is shorter than the number of cycles. Additionally, the fidelity of sequencing by synthesis deteriorates with each additional cycle due to phasing, the desynchronization of cycles that results from unremoved terminator caps ultimately leading to greater uncertainty of base calls in later cycles. Removing adapter contamination and low-quality bases increases the total number of alignable reads, particularly important when analyzing a relatively lower sequence complexity experiment, such as ATAC-STARR-seq. We will use *fastp* to remove sequencing adapters and low-quality reads for the mRNA output and DNA input. A script to perform read trimming can be found here:https://github.com/Bio-protocol/Maize_ATAC_STARR_seq/blob/master/workflow/step01_trim_raw_reads.sh

~~~
# set variables
cd ../
fastqdir=$PWD/FASTQ_files
dna=B73_maize_DNA_input
rna=B73_maize_mRNA_output
threads=16
# trim and filter DNA input reads
fastp -j $dna.json -h $dna.html -w $threads \
   -i $fastqdir/${dna}_1.fastq.gz -I $fastqdir/${dna}_2.fastq.gz \
   -o $fastqdir/${dna}_1.trim.fastq.gz -O $fastqdir/${dna}_2.trim.fastq.gz
# trim and filter mRNA output reads
fastp -j $rna.json -h $rna.html -w $threads \
  -i $fastqdir/${rna}_1.fastq.gz -I $fastqdir/${rna}_2.fastq.gz \
  -o $fastqdir/${rna}_1.trim.fastq.gz -O $fastqdir/${rna}_2.trim.fastq.gzbS - | samtools
sort - > $outdir/B73_maize_mRNA_output.raw.bam
# tidy up log files
mkdir fastp_log_files
mv *.json *.html fastp_log_files
~~~

#### C. Align and process sequenced reads

1. After trimming and quality filtering reads, we align the input DNA and output mRNA reads to the maize B73 version 5 reference genome. To speed up the alignment and downstream processing, we are using 24 CPUs (-t 24) to align the input and output reads. However, users should modify this value to reflect the number of available cores on their system. Additionally, we mark split hits as secondary alignments (-M) to be filtered out downstream as the maize genome is highly repetitive. The output of *bwa mem* is piped to *samtools view* for compression (SAM to BAM) and sorted by alignment coordinate prior to further processing to minimize the footprint on disk. A script to perform these steps can be found here: https://github.com/Bio-protocol/Maize_ATAC_STARR_seq/blob/master/workflow/step02_align_STARR_data.sh

~~~
cd ../
mkdir BAM_files
outdir=$PWD/BAM_files
refdir=$PWD/Genome_Reference
fastqdir=$PWD/FASTQ_files
# align DNA input and pipe to samtools for SAM to BAM conversion
bwa mem -M -t 24 $refdir/Zm-B73-REFERENCE-NAM-5.0.fa
$fastqdir/B73_maize_DNA_input_1.trim.fastq.gz
$fastqdir/B73_maize_DNA_input_2.trim.fastq.gz | samtools view -bS - | samtools sort - >
$outdir/B73_maize_DNA_input.raw.bam
# align RNA output and pipe to samtools for SAM to BAM conversion
bwa mem -M -t 24 $refdir/Zm-B73-REFERENCE-NAM-5.0.fa
$fastqdir/B73_maize_mRNA_output_1.trim.fastq.gz
$fastqdir/B73_maize_mRNA_output_2.trim.fastq.gz | samtools view -bS - | samtools sort - >
$outdir/B73_maize_mRNA_output.raw.bam
~~~
2. To ensure that only high quality alignments remain, here we remove non-properly paired reads (-f 3), secondary hits (-F 256), alignments with low mapping quality (-q 10), and multiple mapped reads (grep -v -E -e ‘\bXA:Z:’) using a combination of *samtools* and unix commands. The header, which contains information on the reference genome and the read mapping parameters, is retained in the output by setting the -h flag in the *samtools view* command.

~~~
# filter DNA input alignments
samtools view -h -q 10 -f 3 -F 256 $outdir/B73_maize_DNA_input.raw.bam | grep -v -E -e
‘\bXA:Z:’ | samtools view -bSh - > $outdir/B73_maize_DNA_input.mq10.pp.unique.bam
# filter mRNA output alignments
samtools view -h -q 10 -f 3 -F 256 $outdir/B73_maize_ mRNA_output.raw.bam | grep -v -E -e
‘\bXA:Z:’ | samtools view -bSh - > $outdir/B73_maize_mRNA_output.mq10.pp.unique.bam
~~~
3. A major difference between analysis of ATAC-seq and STARR-seq data is how assay information is captured by sequencing. For paired-end ATAC-seq, since Tn5 inserts sequencing adapters adjacent to its bound genomic location, chromatin accessibility is encoded as the 5’ ends of sequenced reads. In contrast, STARR-seq produces mRNA transcripts from fragments that are capable of activating their own transcription, thus the entire STARR-seq mRNA and DNA fragment is informative for analysis. The following commands extract the coordinates of sequenced fragments by leveraging the CIGAR strings in BAM paired-end alignments for DNA input and mRNA output. A script to extract fragments can be found here: https://github.com/Bio-protocol/Maize_ATAC_STARR_seq/blob/master/workflow/step03_extract_fragments.sh

~~~
# create output directory
mkdir BED_files
# variables
outdir=$PWD/BAM_files
beddir=$PWD/BED_files
dna=B73_maize_DNA_input
rna=B73_maize_mRNA_output
# extract DNA input fragments (ignore the warnings from bedtools with respect to “missing”
mate pairs, these reflect one of the pairs that had its mate filtered in prior steps)
echo “ extracting fragments from STARR DNA input …”
samtools sort -n $outdir/$dna.mq10.pp.unique.bam \
   | bedtools bamtobed -bedpe -i - \
   | sort -k1,1 -k2,2n - \
   | cut -f1,2,6 - > $beddir/$dna.fragments.bed
# extract mRNA output fragments
echo “ extracting fragments
from STARR mRNA output …”
samtools sort -n $outdir/$rna.mq10.pp.unique.bam \
   | bedtools bamtobed -bedpe -i - \
   | sort -k1,1 -k2,2n - \
   | cut -f1,2,6 - > $beddir/$rna.fragments.bed
~~~

#### D. Identify regions with enriched activity over background

1. To identify regions of the genome with the capacity to activate transcription, we assess enrichment of mRNA reads relative to the input ATAC-seq fragments using *macs2*. As *macs2* is a general peak caller, we need to adjust the default settings to tailor the analysis specifically for ATAC-STARR-seq data. Since this experiment did not use unique molecular identifiers and the number of transcripts from fragment is a direct reflection of its regulatory activity, duplicate mRNA fragments are retained (--keep-dup all). To directly use coverages determined by the input fragment coordinates, we turn off the default fragment shifting model (--nomodel) and set the input type to BEDPE (-f BEDPE). Additionally, we reduce the maximum gap size between candidate peaks to allow for the identification of fine-mapped regulatory elements within a broader regulatory region (--max-gap 50) by setting the minimum peak size to 300 (--min-length 300). Finally, we use the background coverage rates in place of the local bias which aids in peak detection (--nolambda). A script to perform peak calling can be found here: https://github.com/Bio-protocol/Maize_ATAC_STARR_seq/blob/master/workflow/step04_call_peaks.sh

~~~
# prepare output directory and input files
mkdir Peak_data
# generate input files
uniq $beddir/$rna.fragments.bed > $beddir/$rna.fragments.uniq.bed
uniq $beddir/$dna.fragments.bed > $beddir/$dna.fragments.uniq.bed
cat $beddir/$rna.fragments.uniq.bed $beddir/$dna.fragments.uniq.bed | sort -k1,1 -k2,2n - >
$beddir/ALL.fragments.uniq.bed
# find regulatory regions
echo “ calling regulatory regions without duplicate removal …”
macs2 callpeak -t $beddir/$rna.fragments.bed \
  -c $beddir/ALL.fragments.uniq.bed \
  --keep-dup all \
  --max-gap 50 \
  --min-length 300 \
  --nolambda \
  --nomodel \
  -f BEDPE \
  -g 1.6e9 \
  --bdg \
  -n STARR_wdups
# find regulatory regions
echo “ calling regulatory regions with duplicate removal …”
macs2 callpeak -t $beddir/$rna.fragments.uniq.bed \
  -c $beddir/$dna.fragments.uniq.bed \
  --keep-dup all \
  --max-gap 50 \
  --min-length 300 \
  --nolambda \
  --nomodel \
  -f BEDPE \
  -g 1.6e9 \
  --bdg \
  -n STARR_nodups
# find regulatory regions using all unique fragments
echo “ calling regulatory regions by aggregating all unique fragments …”
macs2 callpeak -t $beddir/ALL.fragments.uniq.bed \
  --keep-dup all \
  --max-gap 50 \
  --min-length 300 \
  --nolambda \
  --nomodel \
  -f BEDPE \
  -g 1.69e9 \
  --bdg \
  -n STARR_ALL
# clean-up output
mv STARR_* Peak_data
# merge peaks
cd Peak_data
cat STARR_wdups_peaks.narrowPeak
STARR_nodups_peaks.narrowPeak
STARR_ALL_peaks.narrowPeak | sort -k1,1 -k2,2n - | bedtools merge -i - >
STARR_merged_peaks.bed
~~~
2. To estimate regulatory activity at fine scale, first we need to create a list of unique intervals based on mRNA and DNA fragments. Next, we count the intersection of mRNA and DNA fragments for each unique interval. Finally, we remove all the temporary files to reduce the footprint on disk. A script to estimate enhancer activity can be found here: https://github.com/Bio-protocol/Maize_ATAC_STARR_seq/blob/master/workflow/step05_estimate_enhancer_activity>.sh

~~~
# variables
beddir=$PWD/BED_files
dna=B73_maize_DNA_input
rna=B73_maize_mRNA_output
ref=./Genome_Reference/Zm-B73-REFERENCE-NAM-5.0.fa.fai
# sort the reference
sort -k1,1 -k2,2n $ref > $ref.sorted
# merge RNA/DNA
cat $beddir/$rna.fragments.uniq.bed $beddir/$dna.fragments.uniq.bed \
          | sort -k1,1 -k2,2n - \
          | bedtools genomecov -i - -bga -g $ref.sorted \
          | sort -k1,1 -k2,2n - \
          | cut -f1-3 - > $beddir/Unique_genomic_intervals.bed
# count fragments
bedtools intersect -a $beddir/Unique_genomic_intervals.bed \
          -b $beddir/$rna.fragments.bed \
          -c -sorted -g $ref.sorted > $beddir/$rna.activity.raw.bed
bedtools intersect -a $beddir/$rna.activity.raw.bed \
          -b $beddir/$dna.fragments.bed \
          -c -sorted -g $ref.sorted > $beddir/B73_maize_mRNA_DNA.activity.raw.bed
# clean up temporary files
rm $beddir/Unique_genomic_intervals.bed
rm $beddir/$rna.activity.raw.bed
~~~
3. We and others define enhancer activity as the enrichment of mRNA transcripts that are produced by DNA fragments in the assay in terms of log2(mRNA/DNA) at unique fragment intervals. Prior to taking the log_2_ ratio of mRNA to DNA, we normalize both the input and output to per million to account for differences in sequencing depth and complexity. A pseudocount of one is added to any interval with at least one RNA or DNA fragment. The following code can also be run from the command line using *>Rscript Estimate_Enhancer_Activity*.*R* with the following script: https://github.com/Bio-protocol/Maize_ATAC_STARR_seq/blob/master/workflow/bin/Estimate_Enhancer_Activity.R

~~~
# open an interactive R session to estimate enhancer activity
cd $beddir
R
# load data
a <- read.table(“B73_maize_mRNA_DNA.activity.raw.bed”)
# reformat
rownames(a) <- paste(a$V1,a$V2,a$V3,sep=“_”)
a[,1:3] <- NULL
a <- as.matrix(a)
colnames(a) <- c(“mRNA”, “DNA”)
a <- a[rowSums(a)!=0,]
a <- a + 1
# normalize
a <- a %*% diag(x=1e6/colSums(a))
colnames(a) <- c(“mRNA”, “DNA”)
a <- as.data.frame(a)
# estimate enhancer activity
a$enhancer_activity <- log2(a$mRNA/a$DNA)
# reformat output
rownames(a) <- gsub(“scaf_”,”scaf”, rownames(a))
df <- data.frame(do.call(rbind, strsplit(rownames(a), “_”)))
df$X1 <- gsub(“scaf”, “scaf_”, as.character(df$X1))
mrna <- df
dna <- df
df$X4 <- a$enhancer_activity
mrna$X4 <- a$mRNA
dna$X4 <- a$DNA
# cap negative activity at 0
df$X4 <- ifelse(df$X4 < 0, 0, df$X4)
# save enhancer activity BEDGRAPH file to disk
write.table(df, file=“B73_maize.enhancer_activity.bdg”,quote=F, row.names=F, col.names=F, sep=“\t”)
write.table(mrna, file=“B73_maize.mRNA.bdg”,quote=F, row.names=F, col.names=F, sep=“\t”) write.table(dna, file=“B73_maize.DNA.bdg”,quote=F, row.names=F, col.names=F, sep=“\t”)
# exit interactive mode
q()
~~~
4. To visualize enhancer activity and the normalized mRNA and DNA fragments at any given locus, per million coverage values in bedGraph format from the previous step can be converted into bigwig files (using bedGraphToBigWig from UCSC Utils) for facile visualization using the Integrated Genomics Viewer (IGV) or JBrowse instances.

~~~
bedGraphToBigWig B73_maize.enhancer_activity.bdg ../Genome_Reference/Zm-B73-REFERENCE-NAM-5.0.fa.fai.sorted B73_maize.enhancer_activity.bw
bedGraphToBigWig B73_maize.mRNA.bdg ../Genome_Reference/Zm-B73-REFERENCE-NAM-5.0.fa.fai.sorted B73_maize.mRNA.bw
bedGraphToBigWig B73_maize.DNA.bdg ../Genome_Reference/Zm-B73-REFERENCE-NAM-5.0.fa.fai.sorted B73_maize.DNA.bw
~~~
5. To view the enhancer activity, mRNA, and DNA fragment bigwig files, download and install IGV (https://software.broadinstitute.org/software/igv/download) on your local machine.
6. Unpack, bgzip, and index the gene product annotation. Then load all bigwig, narrowPeak, and genome annotation files using “File > Load from File…” in IGV. An example of an IGV screenshot is shown in **Figure 3**.

~~~
# change directory
cd ../Genome_Reference
# unzip
gunzip Zm-B73-REFERENCE-NAM-5.0_Zm00001eb.1.gff3.gz
# sort by coordinate and remove whole chromosome intervals
sort -k1,1 -k4,4n Zm-B73-REFERENCE-NAM-5.0_Zm00001eb.1.gff3 | grep -v assembly - > Zm-B73-REFERENCE-NAM-5.0_Zm00001eb.1.sorted.gff3
# compress with bgzip
bgzip Zm-B73-REFERENCE-NAM-5.0_Zm00001eb.1.sorted.gff3
# index with tabix
tabix -p gff Zm-B73-REFERENCE-NAM-5.0_Zm00001eb.1.sorted.gff3.gz
~~~

**Figure 3.**
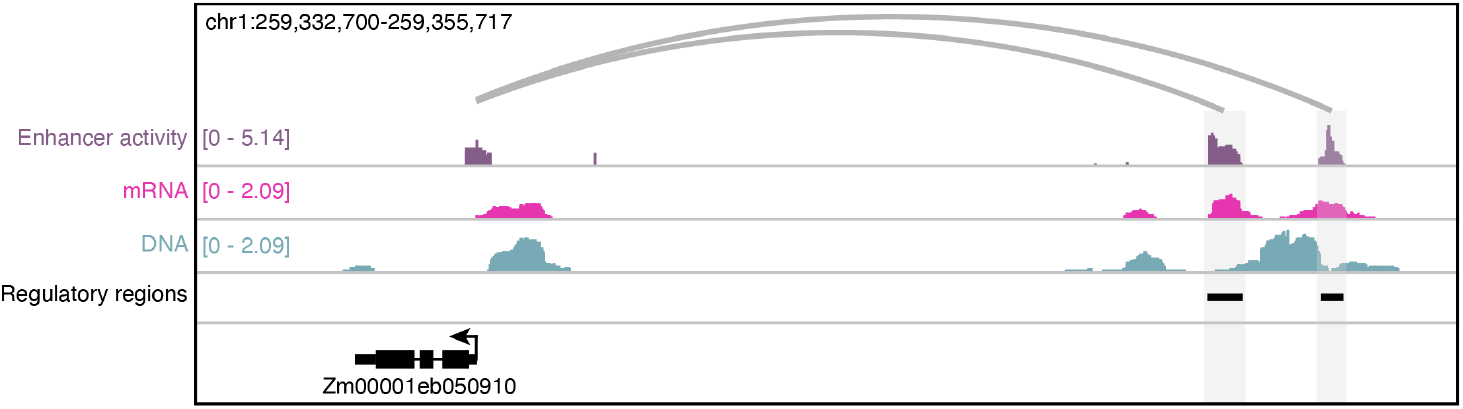
Visualization of ATAC-STARR-seq data in *Zea mays*. Normalized (reads per million) coverages of the DNA ATAC-seq input (blue), self-transcribed mRNA fragments (pink), and enhancer activity (purple; log_2_[mRNA/DNA]) of a 23-kb window. Regulatory regions are shown as black bars, while the grey loops reflect predicted enhancer-gene links.

#### E. Create a list of control regions

To assess enrichment of STARR regulatory regions determined by MACS2 relative to random control regions, we first need to identify regions of the genome that can be uniquely mapped given the sequencing output and read lengths. Although there are numerous methods for estimating mappability, I illustrate a simple approach using synthetic reads tailored to the sequencing parameters of the present experiment. First, we generate the same number of random read pairs with the same sequencing length (36 nucleotides) for mRNA and DNA input using the *wgsim* tool that is supplied to *SAMtools*. The synthetic reads are then remapped and uniformly processed as the original STARR-seq sequencing experiments to identify regions that are uniquely mappable. By constraining randomized control region selection to uniquely mappable genomic intervals, we ensure that downstream comparisons will not be biased by mappability and repeat composition. A script to construct control regions can be found here: https://github.com/Bio-protocol/Maize_ATAC_STARR_seq/blob/master/workflow/step06_create_control_regions.sh

~~~
# move into the “Genome_Reference” directory
cd ./Genome_Reference
# estimate read counts
mRNA_counts=$(samtools view -c ./BAM_files/B73_maize_mRNA_output.raw.bam) DNA_counts=$(samtools view -c ./BAM_files/B73_maize_DNA_input.raw.bam)
# generate simulated reads matching the mRNA output using wgsim from the SAMtools package
wgsim -1 36 -2 36 -d 300 -N $mRNA_counts Zm-B73-REFERENCE-NAM-5.0.fa simulated_STARR_mRNA_r1.fq simualted_STARR_mRNA_r2.fq
# generate simulated reads matching the DNA input
wgsim -1 36 -2 36 -d 300 -N $DNA_counts Zm-B73-REFERENCE-NAM-5.0.fa
simulated_STARR_DNA_r1.fq simualted_STARR_DNA_r2.fq
# compress synthetic fastq files
pigz *.fq
~~~

1. Remap the synthetic reads using the same pipeline as for the original STARR-seq data.

~~~
# variables
outdir=$PWD/BAM_files
refdir=$PWD/Genome_Reference
ref=$refdir/Zm-B73-REFERENCE-NAM-5.0.fa.fai.sorted
fastqdir=$refdir
# align synthetic mRNA output and pipe to samtools for SAM to BAM conversion
bwa mem -M -t 24 $refdir/Zm-B73-REFERENCE-NAM-5.0.fa \
      $fastqdir/simulated_STARR_mRNA_r1.fq.gz \
      $fastqdir/simualted_STARR_mRNA_r2.fq.gz \
      | samtools view -bS - \
      | samtools sort - > $outdir/simulated_STARR_mRNA.raw.bam
# align synthetic DNA input and pipe to samtools for SAM to BAM conversion
bwa mem -M -t 24 $refdir/Zm-B73-REFERENCE-NAM-5.0.fa \
      $fastqdir/simulated_STARR_DNA_r1.fq.gz \
      $fastqdir/simulated_STARR_DNA_r2.fq.gz \
      | samtools view -bS - \
      | samtools sort - > $outdir/simulated_STARR_DNA.raw.bam
~~~
2. Process and filter reads using the original STARR-seq pipeline.

~~~
# filter synthetic mRNA alignments
echo “ filtering synthetic STARR mRNA alignments …”
samtools view -h -q 10 -f 3 $outdir/simulated_STARR_mRNA.raw.bam \
     | grep -v -E -e ‘\bXA:Z:’ \
     | samtools view -bSh - > $outdir/simulated_STARR_mRNA.mq10.pp.unique.bam
# filter synthetic DNA alignments
echo “ filtering STARR DNA alignments …”
samtools view -h -q 10 -f 3 $outdir/simulated_STARR_DNA.raw.bam \
     | grep -v -E -e ‘\bXA:Z:’ \
     | samtools view -bSh - > $outdir/simulated_STARR_DNA.mq10.pp.unique.bam
~~~
3. Identify uniquely mappable regions.

~~~
# extract mRNA output fragments
echo “ extracting fragments from simulated STARR mRNA output …”
samtools sort -n $outdir/simulated_STARR_mRNA.mq10.pp.unique.bam \
     | bedtools bamtobed -bedpe -i - \
     | sort -k1,1 -k2,2n - \
     | cut -f1,2,6 - > $refdir/simulated_STARR_mRNA.fragments.bed
# extract DNA input fragments
echo “ extracting fragments from simulated STARR DNA input …”
samtools sort -n $outdir/simulated_STARR_DNA.mq10.pp.unique.bam \
     | bedtools bamtobed -bedpe -i - \
     | sort -k1,1 -k2,2n - \
     | cut -f1,2,6 - > $refdir/simulated_STARR_DNA.fragments.bed
# merge all fragments (sorting by coordinate at this step may take a while)
cat $refdir/simulated_STARR_mRNA.fragments.bed
     $refdir/simulated_STARR_DNA.fragments.bed \
     | sort -k1,1 -k2,2n \
     | bedtools merge -i - > $refdir/mappable_genomic_regions.bed
~~~
4. Construct control regions using only mappable regions and excluding putative regulatory regions output by *MACS2*.

~~~
# create controls
peaks=$PWD/Peak_data/STARR_merged_peaks.bed
bedtools shuffle -i $peaks \
   -g $ref \
   -incl $refdir/mappable_genomic_regions.bed \
   -excl $peaks \
   | sort -k1,1 -k2,2n - > $PWD/Peak_data/STARR_CONTROL.bed
~~~

#### F. Compare enhancer activity

1. Determine enhancer activity for predicted enhancers output by *MACS2* as well as the negative control regions.

~~~
# create directory to contain analysis
cd ../
mkdir 01_Peak_Analysis
cd 01_Peak_Analysis
# map maximum enhancer activity to putative regulatory regions (wdups)
bedtools map -a ../Peak_data/STARR_merged_peaks.bed -
b ../BED_files/B73_maize.enhancer_activity.bdg -o max -c 4 > STARR_merged_peaks.enhancer_activity.bed
# map maximum enhancer activity to control
bedtools map -a ../Peak_data/STARR_CONTROL.bed -
b ../BED_files/B73_maize.enhancer_activity.bdg -o max -c 4 >
STARR_CONTROL.enhancer_activity.bed
~~~
2. To remove regulatory regions with enhancer activity similar to background, we filter STARR regulatory regions using an empirical false discovery rate (eFDR) based on the matched control regions. A user-specified eFDR threshold identifies the enhancer activity value in the control set that removes 1-eFDR percent of control regions. In this example, we set the FDR to a widely used rate of 0.05. The following code performs and plots eFDR filtering and enhancer activity distributions and can be run from the command line using *>Rscript eFDR_Filter_STARR_Peaks*.*R*. Filtering STARR peaks based on eFDR thresholds derived from the control regions is visualized in **Figure 4A**. An R script of the following code can be found here: https://github.com/Bio-protocol/Maize_ATAC_STARR_seq/blob/master/workflow/bin/eFDR_Filter_STARR_Peaks.R

~~~
# start an interactive R session
> R
# load libraries library(scales)
# load data
starr <- read.table(“STARR_merged_peaks.enhancer_activity.bed”)
con <- read.table(“STARR_CONTROL.enhancer_activity.bed”)
# set missing to 0
starr$V4[starr$V4==‘.’] <- 0
con$V4[con$V4==‘.’] <- 0
# convert to numeric
starr$V4 <- as.numeric(as.character(starr$V4))
con$V4 <- as.numeric(as.character(con$V4))
# get empirical thresholds
fdr <- 0.05
threshold <- quantile(con$V4, (1-fdr))
# filter STARR regulatory regions
filtered <- subset(starr, starr$V4 >= threshold)
# estimate fraction of retained regions
frac <- signif(nrow(filtered)/nrow(starr), digits=4)
# set up multipanel plot area
pdf(“Density_eFDR_STARR_Peak_Filtering.pdf”, width=5, height=5)
# plot control/observed enhancer activities for STARR peaks with duplicates
den.starr <- density(starr$V4)
den.con <- density(con$V4)
plot(NA,
    xlab=“Enhancer Activity”,
    ylab=“Density”,
    xlim=c(range(range(den.starr$x), range(den.con $x))),
    ylim=c(range(range(den.starr$y), range(den.con$y))))
  polygon(x=c(min(den.starr$x), den.starr$x, max(den.starr$x)),
    y=c(0, den.starr$y, 0), col=alpha(“darkorchid4”, 0.5), border=NA)
  polygon(x=c(min(den.con$x), den.con$x, max(den.con$x)),
    y=c(0, den.con$y, 0), col=alpha(“grey80”, 0.5), border=NA)
 abline(v=threshold, col=“red”, lty=2)
 mtext(paste0(“STARR peaks pass = “,frac,” (“, nrow(filtered), “/”, nrow(starr),”)”))
legend(“right”, legend=c(“STARR Peaks”, “Control Peaks”, paste0(“eFDR = “, fdr)),
col=c(“darkorchid4”, “grey75”, “red”), border=c(NA, NA, “red”), pch=c(16, 16, NA), lty=c(NA, NA, 2))
# close device
dev.off()
# save filtered STARR regulatory regions
write.table(filtered, file=“STARR_merged_peaks.enhancer_activity.eFDR05.bed”, quote=F, row.names=F, col.names=F, sep=“\t”)
# exit R
q()
~~~
3. We can now assess the relative enhancer activities across all regions for the filtered and unfiltered STARR peaks and controls using *DeepTools* (**Figure 4B**). A script to plot heatmaps via *DeepTools* can be found here: https://github.com/Bio-protocol/Maize_ATAC_STARR_seq/blob/master/workflow/step07_plot_enhancer_activity.sh

~~~
# load function
getmaps(){
      # input
      ina=$1
      id=$2
      dat=../BED_files/*.bw
      # output
      outa=$id.mat.gz
      outm=$id.mat.txt
      fig=$id.pdf
      # parameters
      threads=48
      window=2000
      bin=20
      cols=YlGnBu
      # create matrix
      computeMatrix reference-point --referencePoint center \
         -S $dat \
         -b $window -a $window \
         -R $ina \
         --missingDataAsZero \
         -o $outa \
         --outFileNameMatrix $outm \
         -p $threads --binSize $bin
      # plot heatmap
      plotHeatmap --matrixFile $outa \
            --colorMap YlGnBu \
            -out $id.heatmap.pdf
}
  export -f getmaps
# run for each file
getmaps STARR_merged_peaks.enhancer_activity.bed STARR_peaks
getmaps STARR_merged_peaks.enhancer_activity.eFDR05.bed STARR_peaks_filtered
getmaps STARR_CONTROL.enhancer_activity.bed control_regions
~~~

**Figure 4.**
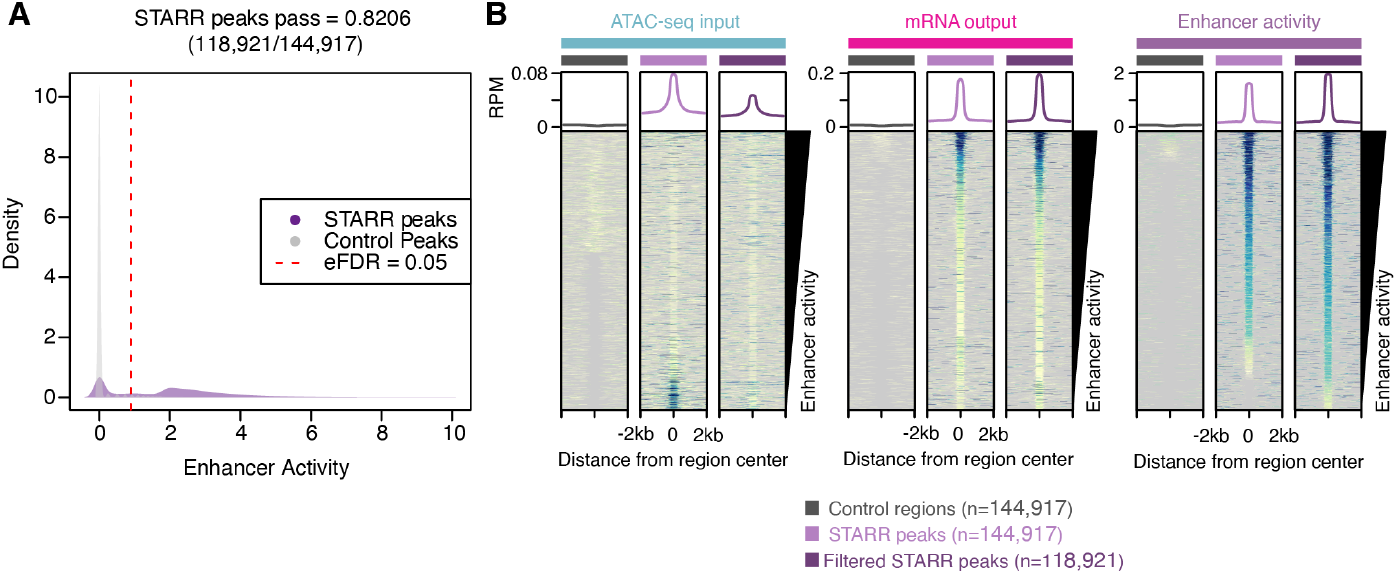
Analysis of STARR regulatory region enhancer activity. **(A)** Distribution of enhancer activity for STARR peaks (purple) and random control regions (grey). Dashed red line indicates the 95% quantile of enhancer activity of random control regions. (**B**) Average (top) and individual site heatmaps of reads per million (RPM) for ATAC-seq input (left), mRNA output (middle) and enhancer activity (right) for control regions, all STARR peaks, and the filtered STARR peak set.

#### G. Identification of large regulatory domains in the maize genome

1. One question these data allow us to ask is whether a relationship exists between the size of a regulatory region and its enhancer activity. So called “super enhancers” in mammalian systems describe hyperactive transcription-activating regulatory domains associated with cell identity that exhibit increased density of TF binding sites compared to typical enhancers (Hnisz *et al*., 2013). Integration of the STARR peaks and enhancer activities with other data sets allows us to determine whether similar hyperactive regulatory domains exist in maize. To query TF binding site density, we first download position weight matrices of known TFs from the *meme* database and identify putative TF binding sites using *fimo* (also from the *meme* suite) conditioning on a *P*-value threshold less than 1e-5. A script to identify large regulatory domains can be found here: https://github.com/Bio-protocol/Maize_ATAC_STARR_seq/blob/master/workflow/step08_identify_large_regulatory_do>mains.sh

~~~
# make a new directory for the TFBS analysis
cd ../
mkdir 02_Hyperactive_Regulatory_Region_Analysis
cd ./02_Hyperactive_Regulatory_Region_Analysis
# download and decompress motif databases
wget https://meme-suite.org/meme/meme-software/Databases/motifs/motif_databases.12.23.tgz
tar -xvzf motif_databases.12.23.tgz
rm motif_databases.12.23.tgz
# variables
threads=16
ref=../Genome_Reference/Zm-B73-REFERENCE-NAM-5.0.fa peaks=../01_Peak_Analysis/STARR_merged_peaks.enhancer_activity.eFDR05.bed controls=../01_Peak_Analysis/STARR_CONTROL.enhancer_activity.bed motifs=./motif_databases/ARABD/ArabidopsisDAPv1.meme
# extract fasta sequences
bedtools getfasta -bed $peaks -fi $ref -fo $peaks.fasta
bedtools getfasta -bed $controls -fi $ref -fo $controls.fasta
# identify putative TFBS
fimo --oc TFBS_peaks $motifs $peaks.fasta
fimo --oc TFBS_controls $motifs $controls.fasta
# reformat fimo output (filtering p-value > 1e-5) using the perl script provided in the github repository (https://github.com/Bio-protocol/Maize_ATAC_STARR_seq/blob/master/workflow/bin/convertMotifCoord.pl)
perl convertMotifCoord.pl TFBS_peaks/fimo.gff | sed -e ‘s/_tnt//g’ - | sort -k1,1 -k2,2n - >
TFBS_peaks.motifs.bed
perl convertMotifCoord.pl TFBS_controls/fimo.gff | sed -e ‘s/_tnt//g’ - | sort -k1,1 -k2,2n - >
TFBS_controls.motifs.bed
# annotate motif coverage/counts for STARR and control peaks
bedtools annotate -
i ../01_Peak_Analysis/STARR_merged_peaks.enhancer_activity.eFDR05.bed -files TFBS_peaks.motifs.bed -both | sort -k1,1 -k2,2n - >
STARR_merged_peaks.enhancer_activity.eFDR05.ann.bed
bedtools annotate -i ../01_Peak_Analysis/STARR_CONTROL.enhancer_activity.bed -files TFBS_controls.motifs.bed -both | sort -k1,1 -k2,2n - >
STARR_CONTROL.enhancer_activity.ann.bed
# extract genes
perl -ne ‘if($_ =∼ /^#/){next;}chomp;my@col=split(“\t”,$_);if($col[2] eq
‘gene’){print”$_\n”;}’ ../Genome_Reference/Zm-B73-REFERENCE-NAM-
5.0_Zm00001eb.1.gff3 | sort -k1,1 -k4,4n - > ../Genome_Reference/Zm-B73-REFERENCE-
NAM-5.0_Zm00001eb.1.genes.gff3
# classify genomic context of STARR and control peaks (you can ignore the warnings from bedtools about inconsistent naming conventions, you can thank the genome assembly team for these annoying, but harmless warnings)
bedtools closest -a STARR_merged_peaks.enhancer_activity.eFDR05.ann.bed -
b ../Genome_Reference/Zm-B73-REFERENCE-NAM-5.0_Zm00001eb.1.genes.gff3 -D b >
STARR_merged_peaks.enhancer_activity.eFDR05.ann2.bed
bedtools closest -a STARR_CONTROL.enhancer_activity.ann.bed -
b ../Genome_Reference/Zm-B73-REFERENCE-NAM-5.0_Zm00001eb.1.genes.gff3 -D b >
STARR_CONTROL.enhancer_activity.ann2.bed
# clean up
mv STARR_merged_peaks.enhancer_activity.eFDR05.ann2.bed STARR_merged_peaks.enhancer_activity.eFDR05.ann.bed
mv STARR_CONTROL.enhancer_activity.ann2.bed
STARR_CONTROL.enhancer_activity.ann.bed
~~~
2. We can now investigate the relationship among regulatory region size, motif density, and enhancer activity to identify putative regulatory domains (**Figure 5A-5F**). To do so, we will start an interactive R session and load the annotated peak and control files from above. A script to automate the following code can be found here: <https://github.com/Bio-protocol/Maize_ATAC_STARR_seq/blob/master/workflow/bin/Characterize_Regulatory_Regio>ns.R

~~~
# open R
> R
# Analyze regulatory regions
# load libraries
library(vioplot)
library(dplyr)
library(MASS)
library(RColorBrewer)
library(scales)
# load data
starr <- read.table(“STARR_merged_peaks.enhancer_activity.eFDR05.ann.bed”)
control <- read.table(“STARR_CONTROL.enhancer_activity.ann.bed”)
# select random control regions to match the filtered STARR peaks
control <- control[sample(nrow(starr)),]
# rename columns for clarity (frac_RR_motif = fraction of regulatory region covered by motifs)
starr[,7:15] <- NULL
control[,7:15] <- NULL
colnames(starr)[4:7] <- c(“activity”, “motif_counts”, “frac_RR_motif”, “gene_distance”) colnames(control)[4:7] <- c(“activity”, “motif_counts”, “frac_RR_motif”, “gene_distance”)
# classify
starr$class <- ifelse((starr$gene_distance < 0 & starr$gene_distance > -200), “TSS”, ifelse(starr$gene_distance < -200 & starr$gene_distance > -2000, “promoter”,
ifelse(starr$gene_distance > 0 & starr$gene_distance < 1000, “TTS”,
ifelse(starr$gene_distance == 0, “genic”, “intergenic”))))
control$class <- ifelse((control$gene_distance < 0 & control$gene_distance > -200), “TSS”, ifelse(control$gene_distance < -200 & control$gene_distance > -2000, “promoter”, ifelse(control$gene_distance > 0 & control$gene_distance < 1000, “TTS”,
ifelse(control$gene_distance == 0, “genic”, “intergenic”))))
# plot distribution
pdf(“STARR_peak_control_genomic_distribution.pdf”, width=10, height=5)
layout(matrix(c(1:2), nrow=1))
pie(table(starr$class))
pie(table(control$class))
dev.off()
# estimate regulatory region size (in log10 scale)
starr$size <- log10(starr$V3-starr$V2)
control$size <- log10(control$V3-control$V2)
# compare sizes between peaks and controls (sanity check)
pdf(“STARR_peak_control_sizes.pdf”, width=5, height=6)
vioplot(starr$size, control$size,
    ylab=“Interval size (log10)”,
    col=c(“dodgerblue”, “grey75”),
    names=c(paste0(“STARR peaks \n (n=“,nrow(starr),”)”),
                  paste0(“Control regions \n (n=“,nrow(control),”)”)))
dev.off()
# compare motif counts
pval <- wilcox.test(starr$motif_counts, control$motif_counts)$p.value
pval <- ifelse(pval==0, 2.2e-16, pval)
mean.peak <- mean(starr$motif_counts)
mean.cont <- mean(control$motif_counts)
# find 95% quantile for control motif count
upper.threshold <- quantile(control$motif_counts, 0.95)
# plot
pdf(“STARR_peak_control_motif_counts.pdf”, width=5, height=6)
vioplot(log1p(starr$motif_counts), log1p(control$motif_counts),
    ylab=“log2(Motif counts + 1)”,
    col=c(“dodgerblue”, “grey75”),
    names=c(paste0(“STARR peaks \n (n=“,nrow(starr),”)”),
           paste0(“Control regions \n (n=“,nrow(control),”)”)),
    ylim=c(0,8),
    areaEqual=T,
    h=0.25)
mtext(paste0(“Wilcoxon Rank Sum P-value = “, signif(pval, digits=3)))
text(1, 7.5, labels=paste0(“Mean = “, signif(mean.peak, digits=3)))
text(2, 7.5, labels=paste0(“Mean = “, signif(mean.cont, digits=3)))
points(1, log1p(upper.threshold), col=“red”, pch=“-”)
points(2, log1p(upper.threshold), col=“red”, pch=“-”)
dev.off()
# split STARR regions by motif counts based on 95% quantile control dist
starr$group <- ifelse(starr$motif_counts >= upper.threshold, “high”, “low”)
pval <- kruskal.test(starr$activity, starr$group)$p.value
pdf(“STARR_peak_activity_vs_group.pdf”, width=5, height=6)
vioplot(starr$activity∼starr$group,
    ylab=“Enhancer activity”,
    col=c(“dodgerblue4”, “dodgerblue”),
    names=c(paste0(“Motif-enriched \n STARR peaks \n (n=“,
nrow(starr[starr$group==“high”,]),”)”),
          paste0(“Motif-depleted \n STARR peaks \n (n=“,
nrow(starr[starr$group==“low”,]),”)”)),
    areaEqual=F,
    xlab=““,
    h=0.25)
mtext(paste0(“Kruskal-Wallis rank sum P-value = “, signif(pval, digits=3)))
dev.off()
# compare STARR region size
pval <- kruskal.test(starr$size, starr$group)$p.value
pval <- ifelse(pval==0, 2.2e-16, pval)
pdf(“STARR_peak_size_vs_group.pdf”, width=5, height=6)
vioplot(starr$size∼starr$group,
   ylab=“Interval size (log10)”,
   col=c(“dodgerblue4”, “dodgerblue”),
   names=c(paste0(“Motif-enriched \n STARR peaks \n (n=“,
nrow(starr[starr$group==“high”,]),”)”),
       paste0(“Motif-depleted \n STARR peaks \n (n=“,
nrow(starr[starr$group==“low”,]),”)”)),
   areaEqual=F,
   xlab=““,
   h=0.25)
mtext(paste0(“Kruskal-Wallis rank sum P-value = “, signif(pval, digits=3)))
dev.off()
# compare motif coverage
pval <- kruskal.test(starr$frac_RR_motif, starr$group)$p.value
pval <- ifelse(pval==0, 2.2e-16, pval)
pdf(“STARR_peak_motif_coverage_vs_group.pdf”, width=5, height=6) vioplot(starr$frac_RR_motif∼starr$group,
    ylab=“Fraction motif coverage”,
    col=c(“dodgerblue4”, “dodgerblue”),
    names=c(paste0(“Motif-enriched \n STARR peaks \n (n=“,
nrow(starr[starr$group==“high”,]),”)”),
           paste0(“Motif-depleted \n STARR peaks \n (n=“,
nrow(starr[starr$group==“low”,]),”)”)),
    areaEqual=F,
    xlab=““)
mtext(paste0(“Kruskal-Wallis rank sum P-value = “, signif(pval, digits=3)))
dev.off()
# split by group
starr.me <- subset(starr, starr$group==“high”)
starr.md <- subset(starr, starr$group==“low”)
write.table(starr.me,
file=“STARR_starrs_peaks.enhancer_activity.eFDR05.ann.high_motif.bed”,
         quote=F, row.names=F, col.names=F, sep=“\t”)
write.table(starr.md,
file=“STARR_starrs_peaks.enhancer_activity.eFDR05.ann.low_motif.bed”,
         quote=F, row.names=F, col.names=F, sep=“\t”)
~~~
3. To determine if the large intergenic regulatory domain regions are associated with cell identity, we will compare enhancer activities versus various cell-type-specific accessible chromatin regions (ACRs) leveraging a recent single-cell ATAC-seq (scATAC-seq) dataset from multiple maize organs (Marand *et al*., 2021). First, download the matrix containing normalized accessibility counts across accessible chromatin regions for each profiled cell type. We then extract ACR genomic coordinates (which are in version 4 of the B73 reference genome) and convert them to version 5 of the B73 reference genome using the *CrossMap* tool and chain file.

~~~
# download the counts matrix
wget -O maize_scATAC_atlas_ACR_celltype_CPM.txt.gz https://www.ncbi.nlm.nih.gov/geo/download/\?acc\=GSE155178\&format\=file\&file\=GSE155178%5Fmaize%5FscATAC%5Fatlas%5FACR%5Fcelltype%5FCPM%2Etxt%2Egz
# unzip
gunzip maize_scATAC_atlas_ACR_celltype_CPM.txt.gz
# download chain file
wget https://download.maizegdb.org/Zm-B73-REFERENCE-NAM-5.0/chain_files/B73_RefGen_v4_to_Zm-B73-REFERENCE-NAM-5.0.chain
# extract coordinates and conform chromosome names to V4 reference
cut -f1 maize_scATAC_atlas_ACR_celltype_CPM.txt \
| grep ‘^chr’ - \
| perl -ne ‘chomp;my@col=split(“_”,$_);print”$col[0]\t$col[1]\t$col[2]\n”;’ - \
| sed -e ‘s/chrB73V4ctg/B73V4_ctg/g’ - \
| sed -e ‘s/chr//g’ - \
| sort -k1,1 -k2,2n - > maize_scATAC_atlas_ACRs.bed
# convert ACR coordinates from V4 to V5
CrossMap.py bed B73_RefGen_v4_to_Zm-B73-REFERENCE-NAM-5.0.chain maize_scATAC_atlas_ACRs.bed > maize_scATAC_atlas_ACRs.V4_to_V5.txt
# discard unmapped and split projections
grep -v ‘Unmap\|split’ maize_scATAC_atlas_ACRs.V4_to_V5.txt \
| perl -ne ‘chomp; my@col=split(“\t”,$_);
print”chr$col[0]_$col[1]_$col[2]\tchr$col[4]_$col[5]_$col[6]\n”;’ - \
| sort -k1,1 -k2,2n - \
| sed -e ‘s/chrscaf/scaf/g’ - \
| sed -e ‘s/chrB73V4_ctg/chrB73V4ctg/g’ - > maize_scATAC_atlas_ACRs.V4_to_V5.clean.txt
# update ACR coordinates in matrix file using R
> R
# read into data frames
conv <- read.table(“maize_scATAC_atlas_ACRs.V4_to_V5.clean.txt”)
mat <- read.table(“maize_scATAC_atlas_ACR_celltype_CPM.txt”)
# subset mat rows by retained ACRs after projection
shared <- intersect(rownames(mat), as.character(conv$V1))
mat <- mat[shared,]
rownames(conv) <- conv$V1
conv <- conv[shared,]
# update mat rowIDs
rownames(mat) <- conv$V2
# save output
write.table(mat, file=“maize_scATAC_atlas_ACR_celltype_CPM.V5.txt”, quote=F,
row.names=T, col.names=T, sep=“\t”)
# exit R
q()
# remove temporary files
rm maize_scATAC_atlas_ACR_celltype_CPM.txt maize_scATAC_atlas_ACRs.bed maize_scATAC_atlas_ACRs.V4_to_V5.txt maize_scATAC_atlas_ACRs.V4_to_V5.clean.txt
~~~
4. Intersect the scATAC-seq ACRs with the STARR peaks with enriched motif counts. Load the scATAC-seq matrix and intersected ACRs/STARR peaks files into R to estimate enhancer activity enrichment scaled by relative accessibilities across cell types. As the STARR-seq data was derived from maize seedlings, we fill further restrict the analysis of scATAC-seq cell types to those derived primarily from maize seedlings. The following code written in R can be executed with the script named ‘motif_enhancer_activity_maize_celltypes.R’ and provides estimates of enhancer activity over various motifs scaled by the relative cell type accessibility, allowing insights into cell-type-specific transcription factor regulation of active enhancers (**Figure 5G**).

~~~
# extract ACR coordinates
cut -f1 maize_scATAC_atlas_ACR_celltype_CPM.V5.txt| grep -v ‘unknown.5.50’ | sed -e
‘s/scaf_/scaf/g’ | perl -ne ‘chomp;my@col=split(“_”,$_);print”$col[0]\t$col[1]\t$col[2]\n”;’ - | sed -
e ‘s/scaf/scaf_/g’ - | sort -k1,1 -k2,2n - > maize_scATAC_atlas_ACRs.V5.bed
# intersect scATAC ACRs with high motif counts STARR peaks
bedtools intersect -a STARR_starrs_peaks.enhancer_activity.eFDR05.ann.high_motif.bed -b maize_scATAC_atlas_ACRs.V5.bed -wa -wb >
STARR_starrs_peaks.enhancer_activity.eFDR05.ann.high_motif.scATAC_ACRs.bed
# map enhancer activity over motifs
bedtools map -a TFBS_peaks.motifs.bed -b ../BED_files/B73_maize.enhancer_activity.bdg -c 4 -o max > TFBS_peaks.motifs.enhancer_activity.bed
# motifs to large regulatory regions
bedtools intersect -a TFBS_peaks.motifs.enhancer_activity.bed -b STARR_starrs_peaks.enhancer_activity.eFDR05.ann.high_motif.scATAC_ACRs.bed -wa -wb
> TFBS_peaks.motifs.enhancer_activity.bed
# open R (alternatively, a script to automate the following code can be found here: <https://github.com/Bio-protocol/Maize_ATAC_STARR_seq/blob/master/workflow/bin/motif_enhancer_activity_maize_>celltypes.R)
> R
# estimate enhancer activity cell type specificity
# load libraries
library(RColorBrewer)
library(gplots)
library(edgeR)
# load data
enh <-
read.table(“STARR_starrs_peaks.enhancer_activity.eFDR05.ann.high_motif.scATAC_ACRs.b ed”)
acrs <- read.table(“maize_scATAC_atlas_ACR_celltype_CPM.V5.txt”)
motifs <- read.table(“TFBS_peaks.motifs.ENRICHED.enhancer_activity.bed”)
# subset for representative leaf-derived clusters
keep <- c(“bulliform.2.26”,
    “bundle_sheath.2.16”,
    “ground_meristem.7.69”,
    “guard_cell.7.74”,
    “guard_mother_cell.7.71”,
    “L1_SAM.4.46”,
    “leaf_provascular.7.67”,
    “mesophyll.2.14”,
    “parenchyma.10.90”,
    “protoderm.7.72”,
    “stomatal_precursor.7.75”,
    “subsidiary.7.68”)
all.acrs <- acrs
# rescale acrs
acrs <- cpm(acrs, log=F)
acrs <- acrs[,keep]
# subset enhancers by genomic feature
enh <- subset(enh, enh$V8==“intergenic”)
# get overlapping regions from the scATAC matrix
enh$ids <- paste(enh$V11,enh$V12,enh$V13,sep=“_”)
enh <- enh[order(enh$V11, decreasing=T),]
enh <- enh[!duplicated(enh$ids),]
shared <- intersect(enh$ids, rownames(acrs))
rownames(enh) <- enh$ids
enh <- enh[shared,]
enh$starrIDs <- paste(enh$V1, enh$V2, enh$V3,sep=“_”)
# filter motifs
motifs$starrIDs <- paste(motifs$V5, motifs$V6, motifs$V7, sep=“_”)
motifs <- motifs[motifs$starrIDs %in% unique(enh$starrIDs),]
# normalize acrs
acrs <- t(apply(acrs, 1, function(x){x/max(x)}))
# iterate over each cell type
cts <- colnames(acrs)
outs <- lapply(cts, function(x){
    access <- acrs[rownames(enh),x]
    names(access) <- enh$starrIDs
    motif.scores <- access[motifs$starrIDs] * as.numeric(as.character(motifs$V15))
    mtf <- data.frame(motif=motifs$V4, score=motif.scores)
    aves <- aggregate(score∼motif, data=mtf, FUN=mean)
    score <- aves$score
    names(score) <- aves$motif
    return(score)
})
outs <- do.call(cbind, outs)
colnames(outs) <- cts
vars <- apply(outs, 1, var)
outs <- outs[vars > 0,]
z <- as.matrix(t(scale(t(outs))))
# # cluster columns
co <- hclust(dist(t(outs)))$order
# reorder rows
z <- z[,co]
row.o <- apply(z, 1, which.max)
z <- z[order(row.o, decreasing=F),]
# cap
z[z < -3] <- -3
z[z > 3] <- 3
# get family
tfs <- data.frame(do.call(rbind, strsplit(rownames(z), “\\.”)))
cols2 <- colorRampPalette(brewer.pal(12, “Paired”))(length(unique(tfs$X1)))
tfs$cols2 <- cols2[factor(tfs$X1)]
# visualize
pdf(“celltype_starr_motif_activity.pdf”, width=10, height=10)
heatmap.2(z, scale=“none”, trace=‘none’,
    RowSideColors=tfs$cols,
    col=colorRampPalette(rev(brewer.pal(9, “RdBu”)))(100),
    useRaster=T, Colv=F, Rowv=F, dendrogram=“none”, margins=c(9,9))
dev.off()
~~~

**Figure 5:**
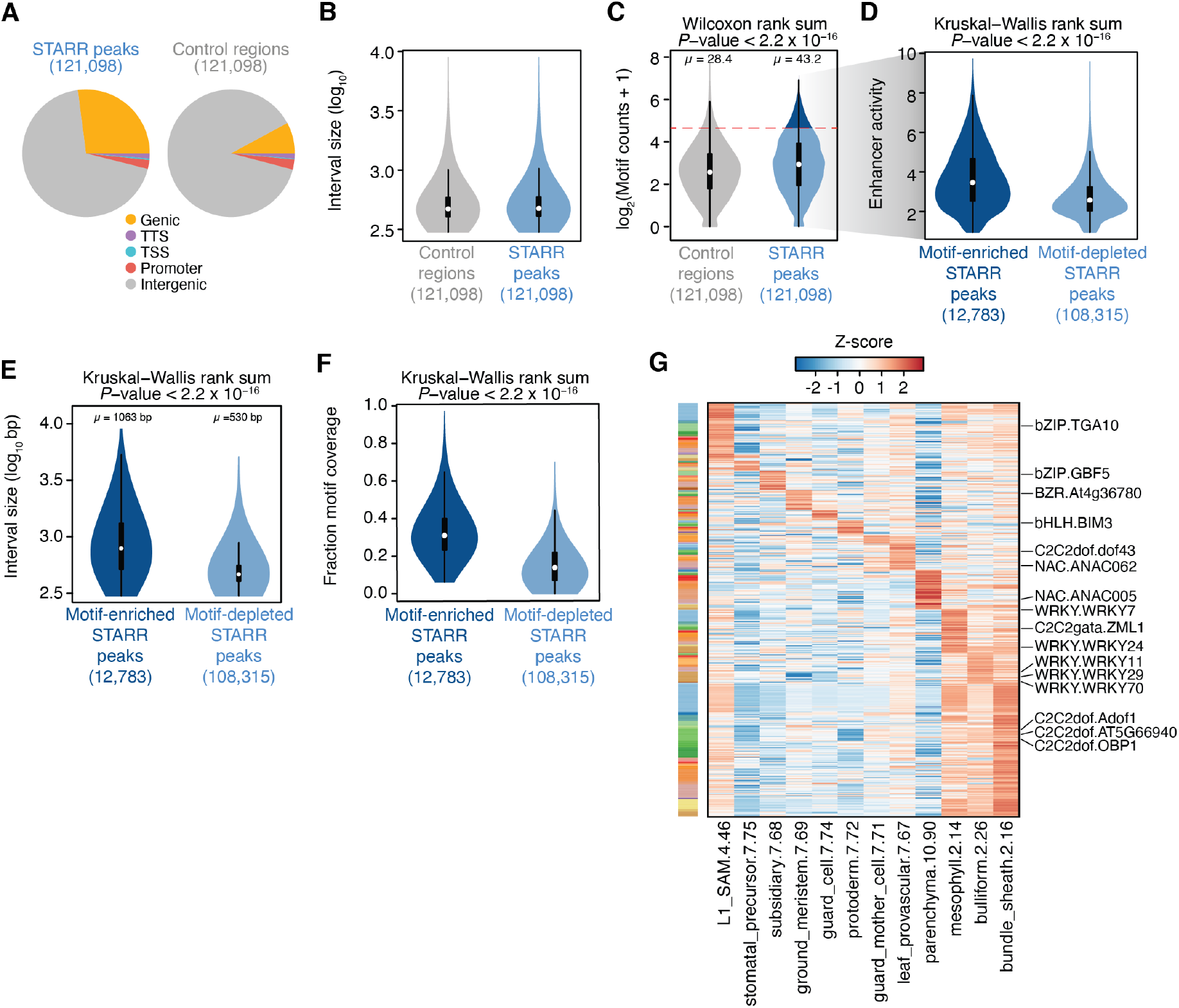
Identification of motif-dense enhancer regulatory domains. (**A**) Genomic distribution of STARR peaks (left) and control regions (right). (**B**) Distribution of control region (grey) and STARR peak (blue) interval lengths. (**C**) Distribution of motif counts in control regions (grey) and STARR peaks (blue). The dashed red line indicates the 95% quantile of motif counts from control regions used to classify STARR peaks into high and low motif count classes. (**D**) Distribution of enhancer activity for STARR peaks with enriched (dark blue) and depleted (light blue) motif counts. (**E**) Distribution of interval lengths for motif-enriched (dark blue) and motif-depleted (light blue) STARR peaks. (**F**) Distribution of fraction of STARR peak covered by motif for motif-enriched (dark blue) and motif-depleted (light blue) STARR peaks. (**G**) Heatmap illustrating Z-score transformed motif enhancer activities across intergenic motif-enriched STARR peaks scaled by the relative chromatin accessibility in various maize cell types.

## Acknowledgments

This study was funded by support from the National Science Foundation (DBI-1906869) and the National Institute of Health (1K99GM144742) to A.P.M. The ATAC-STARR-seq data analyzed in this study was originally generated by Ricci, Lu, Ji and colleagues (Ricci *et al*., 2019).

## Supplementary information

1. Data and code availability: All data and code have been deposited to GitHub: https://github.com/Bio-protocol/Maize_ATAC_STARR_seq

